# Limits of noise for autoregulated gene expression

**DOI:** 10.1101/100115

**Authors:** Peter Czuppon, Peter Pfaffelhuber

## Abstract

Gene expression is influenced by extrinsic noise (involving a fluctuating environment of cellular processes) and intrinsic noise (referring to fluctuations within a cell under constant environment). We study the standard model of gene expression including an (in-)active gene, mRNA and protein. Gene expression is regulated in the sense that the protein feeds back and either represses (negative feedback) or enhances (positive feedback) its production at the stage of transcription. While it is well-known that negative (positive) feedback reduces (increases) intrinsic noise, we give a precise result on the resulting fluctuations in protein numbers. The technique we use is an extension of the Langevin approximation and is an application of a central limit theorem under stochastic averaging for Markov jump processes (Kang, Kurtz and Popovic, 2014). We find that (under our scaling and in equilibrium), negative feedback leads to a reduction in the Fano factor of at most 2, while the noise under positive feedback is potentially unbounded. The fit with simulations is very good and improves on known approximations.

## Introduction

It is now widely accepted that gene expression is a stochastic process. The reason is that a single cell is a system with only one or two copies of each gene and of the order tens for mRNA molecules [1, 2, 3]. Experimentally, this stochasticity can even be observed directly by single-cell measurements such as flow cytometry and fluorescence microscopy, which show the inherent fluctuations of protein numbers arising from cell to cell [4].

Usually, noise in gene expression is divided into an intrinsic and an extrinsic part [1, 5]. While the intrinsic part leads to variation of protein numbers from cell to cell in the same environment, the extrinsic part is attributed to the different environmental conditions of the cell. In practice, ensemble averages eliminate intrinsic noise, while single-cell measurements can be thought of having a constant environment, thus eliminating extrinsic noise [6, 7].

Stochasticity in gene expression is not only interesting per se. Today, its role in evolution, development and cell fate decisions is under discussion [8, 9, 10, 11, 12]. In general, noise should be detrimental to cells, since they have to function constantly. Therefore, mechanisms reducing the level of noise are beneficial for real systems.

Under the central dogma of molecular biology, modeling stochasticity of gene expression is straight-forward (see [13] for a review). A gene, which is either turned on or *off,* is transcribed into mRNA, which is translated into protein. Both, mRNA and protein are degraded at constant rates. Since the resulting chemical reaction network is linear, the master equation can be solved and all moments can be derived analytically. Most interestingly, the variance can be decomposed into the effects of switching the gene on and *off,* noise due to the finite life-time of mRNA, and random fluctuations in the production of protein [13]. It is often stated that gene expression tends to occur in bursts, which occur due to the short life-time of the on-state of the gene and due to the short life-time of mRNA [14].

We are interested in the effect of self-regulation on gene expression noise. It is known that a negative feedback loop, i.e. a protein suppressing its own transcription (or translation) leads to a reduced noise, while positive feedback is attributed to increase in noise ([15, 16]). Although these findings are wide-spread, a complete mathematical analysis is lacking. At least, for negative feedback, [17] and in more generality [18] quantify the effect of negative feedback using a linearization argument. The latter paper further analyzes different feedback models differing between translational and transcriptional autoregulation. Moreover, [19] derive the equilibrium distribution using a multi-scale approach under negative feedback.

Most analyses of noise in unregulated gene expression rely on the master equation (e.g. [13]). By the linearity of this equation, a solution can be given explicitly. Using the approximation that the gene is constantly transcribed to mRNA, this linearity can as well be used under negative feedback [17, 18]. Our approach differs in two ways. First, we are using martingale methods from stochastic analysis in order to describe the chemical system [20]. Second, we can relax the assumption that the gene is transcribed constantly, and therefore derive a more general result. Consequently, we are able to analyze noise in a truly non-linear system under a quasi-steady-state assumption.

While the full model of regulated gene expression (or any other chemical reaction network) is usually hard to study, considering an ODE approach instead, which approximate the full model, leads to new insights. Formally, a law of large numbers – usually referred to as a fluid limit – can be obtained connecting the stochastic and deterministic model [21, 22]. While such a law of large numbers gives a deterministic limit, fluctuations are studied using central limit results; see [23]. The special situation for gene expression is that the gene and mRNA only have a few copies, while the protein is often in large abundance. Such multiscale models are often studied under a quasi-steady-state assumption [24]. Here, the species in low abundance are assumed to evolve fast, such that the slow, abundant, species only sense their time-average. For such a stochastic averaging, not only a law of large numbers is given e.g. by [25], but also a central limit result has recently be obtained by [26].

While a multi-scale approach to stochastic gene expression is not new (see [27, 19]), the analysis of fluctuations for such systems is not finished yet. In a diffusion setting, fluctuations for a multiscale system were computed using a Poisson-equation [28, 29]. The results by [26] are similar but are based on Markov jump processes. As our results show, fluctuations take into account all sources of noise and we give explicit formulas for the reduction of noise under negative feedback and the increase in noise under positive feedback.

## 1 The model

We are dealing with the standard model of gene expression without and with feedback; see e.g. [19]. (All details are found in the SI.) Using the terminology from [13], we write for the model without feedback (or the *neutral* model)

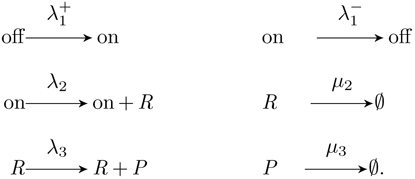

Here, off and on refer to the inactive and the active gene, respectively. mRNA is denoted by *R,* and the protein by *P*. Exchanging the first line by

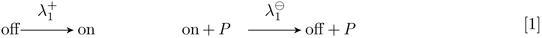

then models a negative feedback while

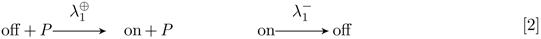

models a positive feedback. (Note that our results will not change if the protein binds to the gene in the sense that on + P → off or off + P → on.) We will refer to these three full models by [*neu*], [*neg*] and [*pos*], respectively. We write *R*(*t*) and *P*(*t*) for the number of mRNA and protein in the system at time *t*.

In the sequel, we are interested in the situation where the gene and RNA has *O*(1) copies, while the protein is much more abundant (i.e. *O*(*N*) for some large *N*). For the parameters in [*neu*], this means that 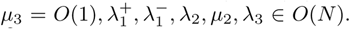. For [*neg*], we have in contrast that 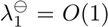 since then 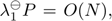, while for [*pos*] we have 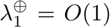 since then 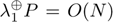. For this scaling, on, off and *R* are *fast* molecular species, and quickly equilibrate. Using a quasi-steady-state assumption, we see that, approximately, (see SI, Theorem 1),

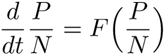

with

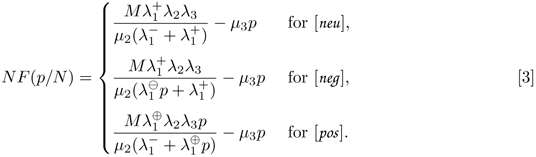

Our goal is derive the variance in protein numbers under [*neu*], [*neg*] and [*pos*]. While [*neu*] is solved explicitly elsewhere, e.g. in [13], some approximations have to be made for [*neg*] and [*pos*]. A first idea is to use a Langevin approximation and write

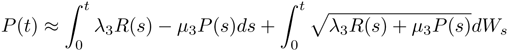

and *v_P_* for the exact solution of 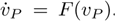. Then, for 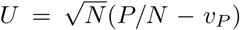 and *b(v_P_*) = λ_3_𝔼_π_ [*R*] + *μ_3_v*_*P*_, the average diffusion for fixed *v_P_*, i.e.

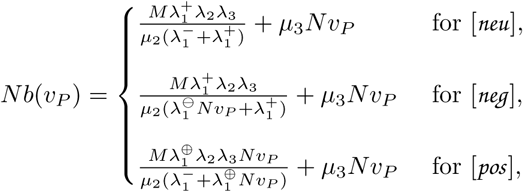

we find

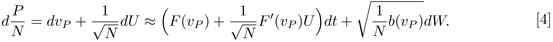

This approach builds on applying a quasi-steady-state approach whenever possible, i.e. when averaging over the on/off-state of genes in order to derive *F*, and the number of RNA, which is approximated by its mean in order to derive *b*. Consequently, fluctuations arising from these two mechanisms cannot be accounted for in the resulting variance. In the next section, we give our results which take all reactions into account for deriving noise in gene expression.

## 2 Results

### Approximate variance and Fano factor for *P*

As an application of [26], we derive in the SI (Theorem 2) the following central limit result: Let *v_P_* be the exact solution of 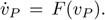. The deviations of *P*/*N* from *v_P_*, given through 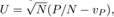, satisfy for large *N* the SDE

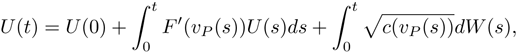

with *W* a one-dimensional standard Brownian motion and

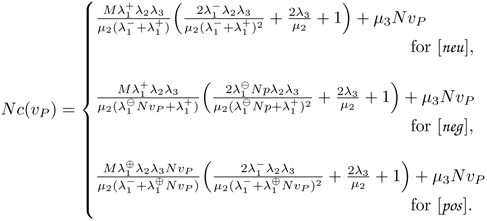

Solving for *P*, we obtain, approximately (compare with [4])

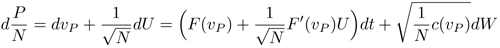

While this limit provides a dynamical result along paths of *P*, we can also use this approximation and study the process in equilibrium by setting 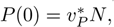 where (see SI, Section B)

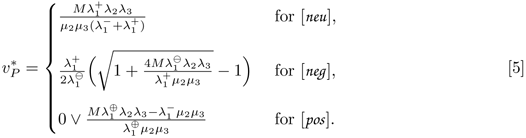

is the unique solution of *F*(*v_P_*) = 0 (and hence 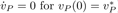). We obtain in this case

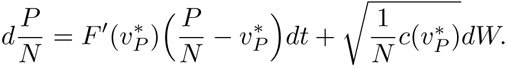

This is an Ornstein-Uhlenbeck process and it is well-known that its equilibrium has the Fano factor

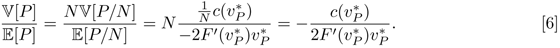

Since no factor *N* appears on the right hand side, some authors call the Fano factor dimensionless. Empirically, it was found e.g. by [30], that for all classes of genes and under all conditions, the expression variance was approximately proportional to the mean, which is again reminiscent of the lacking *N* in the Fano factor above. Plugging in the quantities from above, we find with

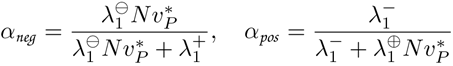

that (see SI, Remark B.1)

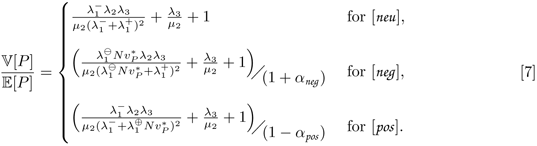

### Refining the Fano factor

Using a slightly different model where *μ*_2_ = *O*(1) (hence *R* = *O*(*N*)) and λ_3_ = *O*(1) (hence still *P* = *O*(*N*)), it is possible to derive a refined formula for the Fano factor. In SI, Section C, we derive (see (SI.13), (SI.15) and (SI.16)) that

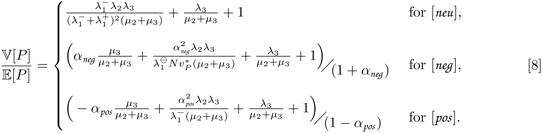

Note that this equation approximately gives [7] for μ_2_, λ_3_ >> 1. In practice (and in our simulations below), the life-time of proteins is much larger than the life-time of mRNA, such that [8] doesn’t produce a better fit than [7]. Therefore, we will use [7] in the sequel.

### Comparing the noise in [*neu*], [*neg*] and [*pos*]

It is frequently reported that a negative feedback in gene expression results in a reduced variance (noise) of protein levels, whereas a positive feedback enhances noise. These observations can be made precise by our results from above. Here, we report some consequences on the equilibrium variance and the Fano factor, 𝕍[*P*]/𝔼[*P*]. For a fair comparison, we use the models [*neu*], [*neg*] and [*pos*] for equal values of 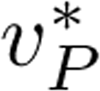. More specifically, we use parameters which satisfy 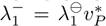 for [*neg*] and 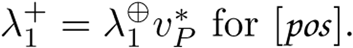 Then, from [7], we find that (see SI, Section B.2)

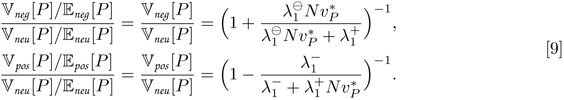

**Figure 1:**
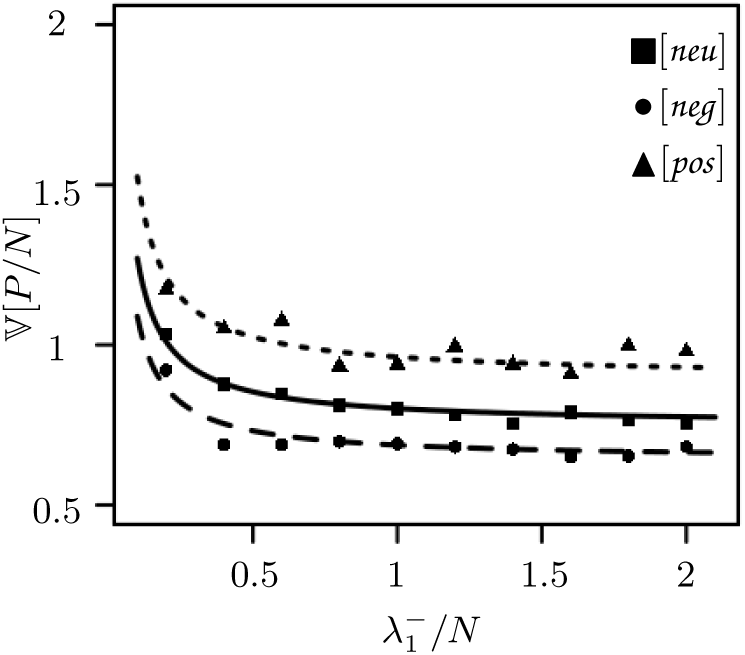
Simulations and theoretical results with a fixed mean of 1250 proteins. The gene association and dissociation rates are varied, i.e. 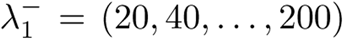 and 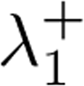 is chosen such that the protein mean equals 1250 in each case. Furthermore, these rates are adjusted in the cases of negative and positive feedback according to the description right before [9] for the different cases. The dissociation rate 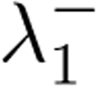 is plotted on the x-axis whereas the y-axis represents the variance in protein numbers. The solid, dotted, dashed line are the neutral, positive and negative case, respectively. The other parameter are given by *M* = 1,*N* = 100, λ_2_ = 300, μ_2_ = 100, λ_3_ = 500 and *μ_3_* = 1. Each data point is derived from 1000 Monte-Carlos simulations (cf. [31]) of the full system given by [*neu*], [*neg*] and [*pos*].

In particular, we see that the variance is reduced in [*neg*] and increased in [*pos*], as expected. Furthermore, we performed simulations and compared them to our predictions. As can be seen in Figure 1 the simulated results for relatively small *N*, i.e. small numbers of proteins, fit the predictions quite well. Of course this fit improves by increasing values of *N*.

### Comparison to previous results

Here, we compare our results in the neutral case with [13], and with [17], [19] and [18] in the case of negative feedback.

*The neutral case:* Using the approximation *μ_3_ << μ_1_, μ_2_*, i.e. the protein life-time is much longer than the duration of constant gene activity and RNA life, we see in the SI, Section D.1, that (4) from [13] and [7] produce the same results.

*Negative Feedback, [17]:* For [*neg*], a linearization was studied in [17] for the case of fast switching on and off of the gene. (This will mean that both rates for switching on and off the gene are large; see SI, Section D.2.) Using the further approximation that the gene is switched on most of the time,

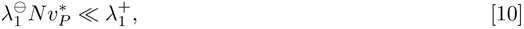

they find the approximation

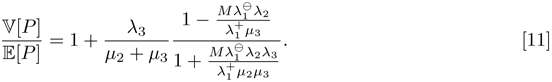

Using that *μ_3_ << μ_2_* and [10], we can rearrange this to give

**Figure 2:**
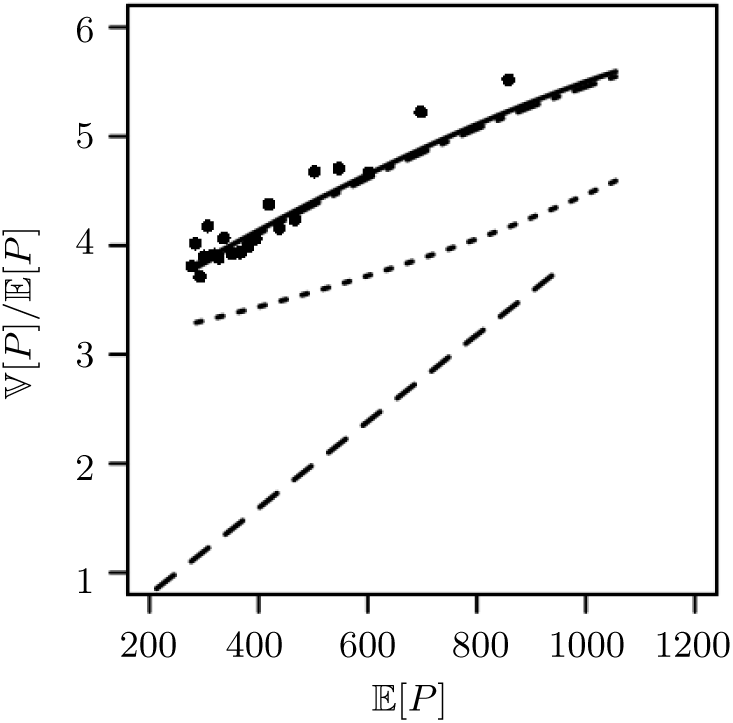
Simulations and theoretical results of gene expression with negative feedback. The mean given on the x-axis is varied and plotted against the Fano factor on the y-axis. The solid line represents [7], the dash-dotted line the result in [8], the dashed line the result from [17] given in [11] and the dotted line the Fano factor calculated in [18] given in [12]. The parameter for the simulations are chosen as follows: *M* = 1,*N* = 100, 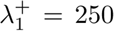, 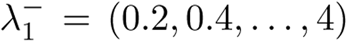,λ_2_ = 300, *μ*_2_ = 100, λ_3_ = 500, *μ_3_ = 1*. The bullets represent the estimated Fano factors of the full system [*neg*] obtained from 1000 Monte-Carlo simulations (cf. [31]) for each value of 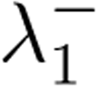.

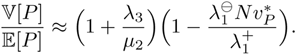

Since fluctuations of switching the gene on and off are not accounted for in this calculation, we therefore find that, in this approximation,

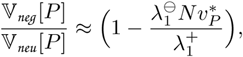

which approximately equals [9]. Therefore, the results of [17] are similar to our results, but less accurate if gene (in-)activation is slower or if [10] is not satisfied; see also Figure 2.

*Negative Feedback, [18]:* As explained around [4], the usual Langevin approximation cannot account for all fluctuations when a quasi-steady-state assumption is made. (Precisely, it cannot account for fluctuations in the averaged variables.) In [18], a Langevin approach is carried out in order to analyze fluctuations in autoregulatory gene expression in the cases of transcriptional and translational feedback. The author considers the mRNA and the protein to evolve on the same time-scale whereas the gene (or DNA) is considered to be on a faster time-scale. For transcriptional feedback (which we study here), he obtains in his (5) –see SI, Section D.3 for the transformation of their results into our parameters

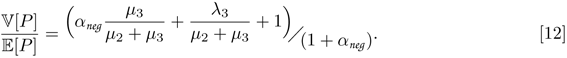

This corresponds exactly to [8] with a missing term in the numerator. This term arises from fluctuations in gene activation, which was averaged out in calculations done [18]. At least, [12] arises from [8] if we assume that 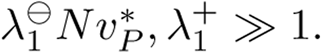.

*Negative Feedback, [19]:* Our result for [*neg*] has Theorem 2 of [19] as a limit result. They study a similar model (with *M* = 1), but with a scaling such that λ_3_*,μ_3_* = *O*(1) and 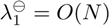, leading to low (i.e. *O*(1)) abundance of protein in the system. The resulting birth-death process for *P* has a stationary distribution which they compute explicitly. Moreover, setting

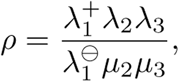

they obtain in their Corollary 4.1 that, in equilibrium,

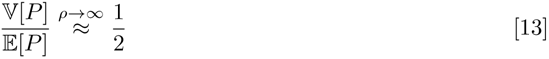

For large ρ, it is 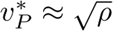 and [7] gives the same limit.

### Simpler model of gene expression

In the literature simpler models of gene expression are studied as well. Here, only two molecular species are involved. Either, the gene is constitutively expressed, therefore ignoring the state of the gene, or translation is neglected and the gene is assumed to be transcribed leading to protein in one step [32, 33, 34]. In either case, we have the model

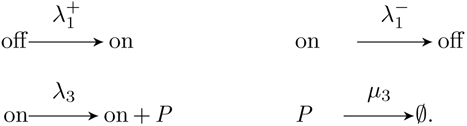

for [*neu*], whereas for [*neg*] and [*pos*], we take [1] and [2] instead of the first line, respectively. We note that this model arises from the full model described above (see SI, Section D.4), when letting *μ_2_* = λ_2_ → ∞. Hence, we obtain for the simpler model the approximation

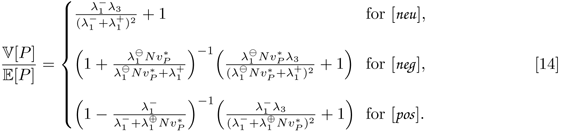

## 3 Discussion

Quantifying noise in gene expression is essential for understanding regulatory networks in cells [17]. Our results give the most complete theory on the intrinsic noise available today. While negative feedback is known to reduce noise under negative feedback, we improve on the quantification of this effect. Moreover, we can provide the same quantification also for positive feedback, where noise is increased. In particular, [9] shows that the average time the gene is *off* determines the reduction of noise in all cases relative to unregulated genes; see also [35]. Both, for negative and positive feedback, the change in noise is maximal if the gene is *off* most of the time, while still having the same amount of protein as in the unregulated (neutral) case. This finding is reminiscent of the fact that gene expression comes in bursts. The burstiness is most extreme if the gene is *on* only for a short time, producing a large amount of mRNA, and afterwards *off* for a long period. Interestingly, previous approaches only gave approximations for noise for negative feedback if switching the gene on and off is very fast [17, 18] and if the gene is *on* most of the time [17]. Hence, all previous papers could not have seen the effects of gene (in-)activation on protein noise. As in previous results [19], we find that in the limit where the gene is *off* most of the time, the negative feedback reduces noise at most by a factor of two. In addition, noise can increase unboundedly for positive feedback.

Today, quasi-steady-state assumptions are frequently used when analyzing chemical reaction networks. While the intuition suggests the correct approach when approximating the system by a deterministic path, studying fluctuations is apparently much less obvious. In [36], some special cases are studied when a straight-forward approximation of the fluctuations work. In our analysis, we use a new approach by [26] and can also interpret all terms arising in [7]. E.g., for negative feedback, we find – as in the neutral case – contributions from randomness in translation and transcription by

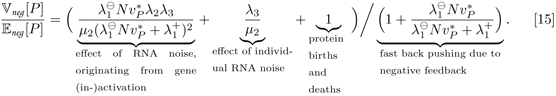

Moreover, the negative feedback pushes the amount of protein faster back to its equilibrium value for a burst of gene expression. This results in the denominator in [15], which has the biggest< effect of the noise-reducing effect of negative feedback. In addition, another source of noise comes from switching the gene on and off (term 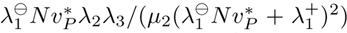). It is due to the latter term that the fit of simulations and theory (see e.g. Figure 2) is excellent. Previous studies have averaged out this source of noise, and only the recent approach of [26] reveals the impact of switching gene on and off on the noise in protein numbers.

In their paper, [26] gave as an example an approximation of noise for Michaelis-Menten kinetics and a model for virus infection. Their method relies mostly on solving a Poisson equation *L*_2_*h* = *F_n_ — F*, where *L*_2_ is the generator of the fast subsystem (gene and RNA in our example), *F_N_* and *F* describe the evolution of the slow system (protein) including all fluctuations and in the limit using the quasi-steady-state assumption, respectively. We stress that this approach is not only useful for equilibrium situations, but also for understanding noise if the slow system has not reached equilibrium yet, e.g. after a cell split.

It was argued that complexity of gene regulatory networks leads to a reduction in the level of noise, while certain network motifs always lead to increased levels of noise [37, 38]. Experimentally, gene expression noise can be used to understand the dynamics of gene regulation [39]. Our analysis should provide an approach for distinguishing between different models of gene regulation based on measurements of noise levels.

## Authors’ contributions

PC and PP conducted the research and wrote the manuscript.

## Acknowledgements

We thank Jens Timmer, Freiburg, for discussion.

## Supporting Information

## A The model

We are dealing with gene expression without and with transcriptional feedback; see e.g. Swain (2004); Dessalles et al. (2016). For general formulations of chemical reaction networks (and their mathematical representations), consult Anderson and Kurtz (2015). Using the terminology from Paulsson (2005), we write for the model without feedback (or the ***neutral*** model)

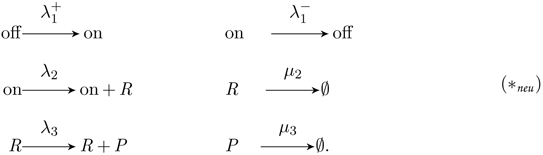

Here, *off* and *on* refer to an inactive and an active gene, respectively. The mRNA is given by *R*, and the protein by *P*. While the first line of chemical reactions models gene switching from *off* to *on* and back, the second line encodes transcription and degradation of mRNA, while the third line gives translation and degradation of proteins. Exchanging the first line by

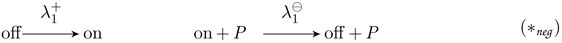

then models a negative feedback while

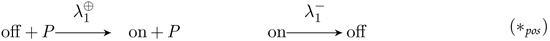

models a positive feedback. In all cases, we number the equations from left to right and from top to bottom by 1–6, so *K* = {1,…, 6} is the set of chemical reactions. The species counts are given by *X*_off_, *X*_on_,*X*_R_ and *X*_P_ for inactive and active gene, mRNA and protein, respectively. We will make use of the following scaling for the abundances of chemical species

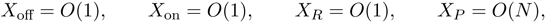

or

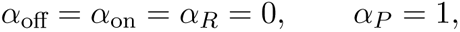

such that *X_i_* = *O*(*N*^α*i*^) for all *i*. Reactions are scaled for all models by

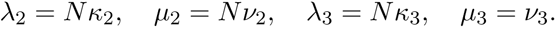

For the neutral model, we also set

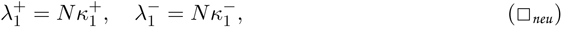

whereas for negative feedback

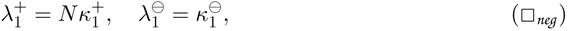

and for positive feedback

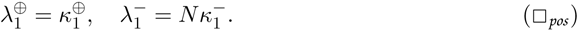

(Note that this scaling obeys 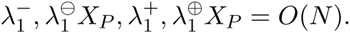.) Setting 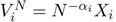, and for *M* copies of the gene, we have in the neutral case

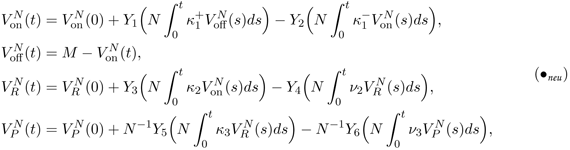

for independent, rate 1 Poisson processes *Y*_1_,…,*Y*_6_. The first equation changes in the case of negative feedback to

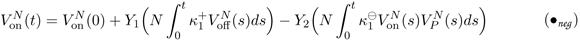

and in the case of positive feedback to

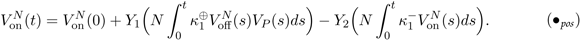

In the sequel, we will refer to the model without, negative and positive feedback simply as [*neu*], [*neg*] and [*pos*], respectively. We understand all equations (*_*neu*_), (□ _*neu*_), (●_*neg*_) as the bases for [*neu*], equations (*_neg_), (□ _*neg*_), (●_*neg*_) as the bases for [*neg*] and all equations (*_*pos*_), (□ _*pos*_), (●_*pos*_) as the bases for [*pos*].

## B Results

### B.1 Law of Large Numbers and Central Limit Theorem

The following result can be obtained using a quasi-steady-state assumption. It relies on the method of stochastic averaging; see e.g. Ball et al. (2006).

#### Theorem 1 (Law of Large Numbers)

*We consider the models [neu], [neg] and [pos] and assume that* 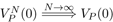 *Then,* 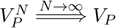, where *V*_P_ solves

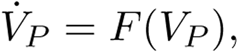

*with (see also [3] in the main text)*

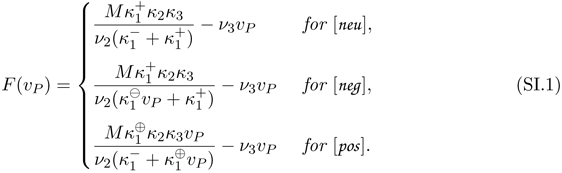

*In particular, the equilibrium is given by (see also [5] in the main text)*

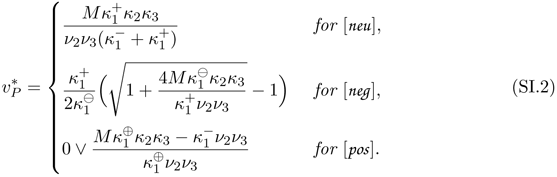

*Proof.* The equilibrium π on the fast scale satisfies, for *V_P_* = *v_P_* fixed,

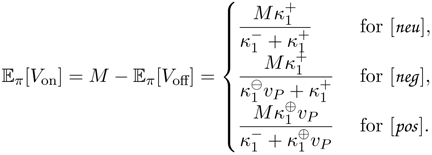

and

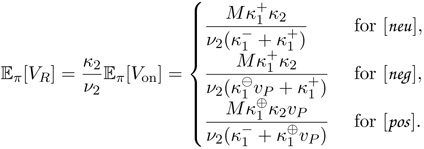

Plugging this equilibrium into the equations for *V_P_*, we obtain that

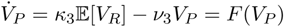

with *F* as in (SI.1). Computation of the equilibria is standard by solving *F*(*v_P_*) = 0. In particular, we have to solve

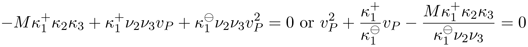

for the equilibrium of [*neg*].□

Our next goal is to show that 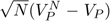 converges and to determine the limiting process. In the proof, we will make use of the method developed by Kang et al. (2014), which we recall in Section E.

#### Theorem 2 (Central Limit Theorem)

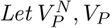, *and F be as in Theorem 1 and* 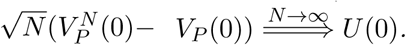*Then, for the models* [*neu*], [*neg*] *and* [*pos*], 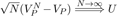 *where U solves*

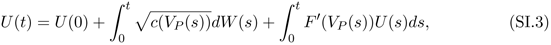

*with W the one-dimensional standard Brownian motion and*

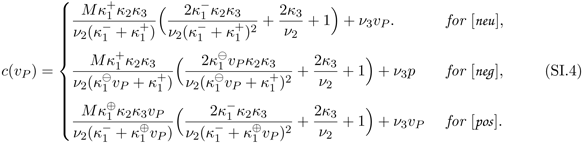

**Remark B.1** (Deriving the Fano factor in equilibrium). In order to compute the approximate variance of 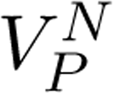, when started in the equilibrium 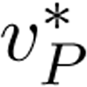 we make use of the fact that the SDE (SI.3) is solved by an Ornstein Uhlenbeck process. In particular, we obtain at late times (see also [6] in the main text)

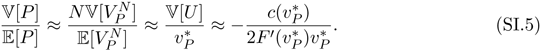

In order to compute the right hand side, note that

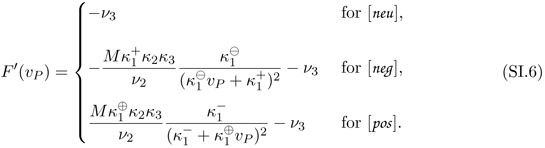

Plugging in the equilibrium 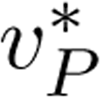 from (SI.2) or [5] for [*neg*] and [*pos*], we obtain in particular that

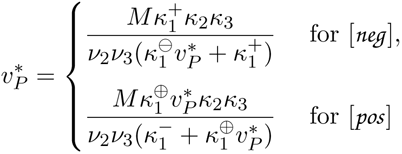

and therefore

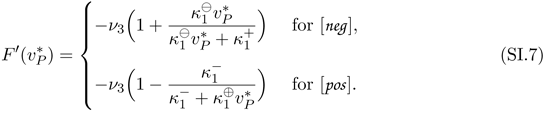

In addition,

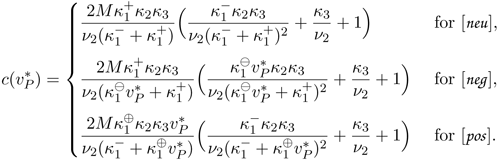

Hence, plugging these quantities into (SI.5) gives

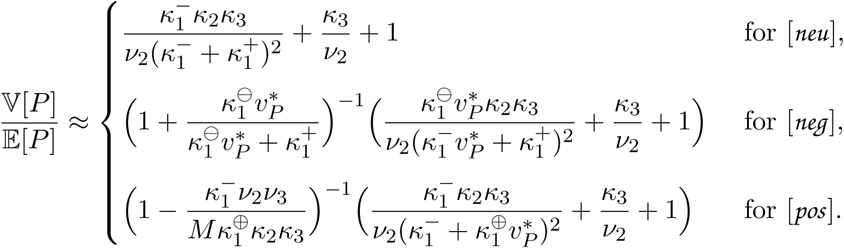

*Proof of Theorem 2.* We have to show (➊)-(➍) from Section E in all cases. Note that the function *F* from Theorem 1 already satisfies (➊). In all cases, the system (*V*_on_, *V*_R_, *V*_P_) is a Markov process with a generator of the form (SI.21) with

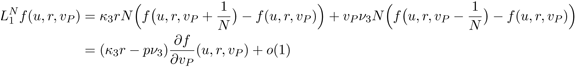

(and different operators 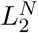). This already implies that for all cases

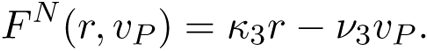

For [*neu*],

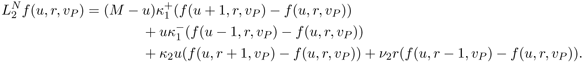

From (➋) and (SI.1), we see that we need to solve

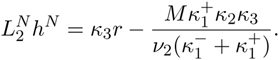

Choosing the Ansatz

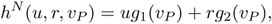

we obtain that

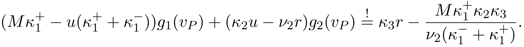

Solving for *g*_2_ and then for *g*_1_, we obtain

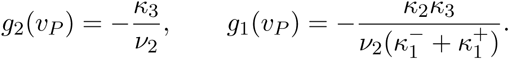

Then, 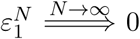 since *h^N^* is bounded in *N* and 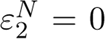 by construction. Hence, we have shown (➌). For (➍), if π is the equilibrium of the fast species *U*, *R* for given value *v_P_* of the slow species as in Theorem 1, we have that (using notation introduced in Section E)

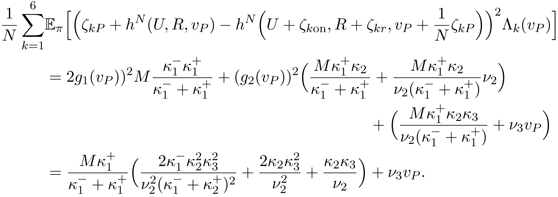

For [*neg*], all calculations above are the same, but with 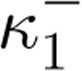 replaced by 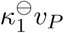, and for [*pos*], all calculations are the same with 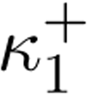 replaced by 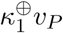.□

### B.2 Comparing [*neu*], [*neg*] and [*pos*] if 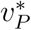 is equal

Consider a model [*neu*] with parameters 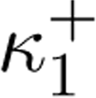, 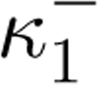, κ_2_, κ_3_, ν_2_, ν_3_ and let 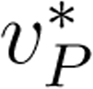 be the equilibrium from [5] or (SI.2). In addition, consider a model [*neg*] with 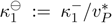 and all other parameters as above and a model [*pos*] with 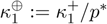 and all other parameters as above. Then, from (SI.1), we see that all models have 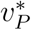 as their unique deterministic limit with the same

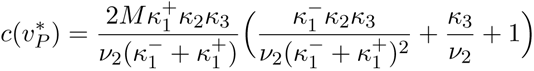

from (SI.4). Plugging all quantities in (SI.5) (or [6]) then gives

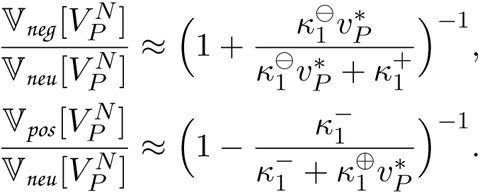

## C Slow RNA

### C.1 The model

Here, we study the case

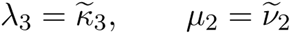

which leads to *X_R_* = *O*(*N*), such that we have the scaling α_R_ = 1 or *V_R_* = *X_R_*/*N*. Hence, *V*_on_ is fast and (*V_R_*, *V_P_*) are slow. Note that in this case, we have that 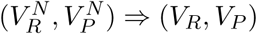 with

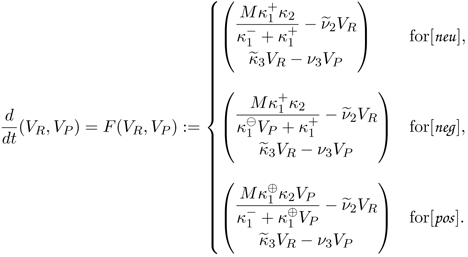

#### Theorem 3 (Central Limit Theorem)

*Let* 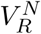, *V_R_*, 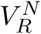, *V_p_ and F be as above and* 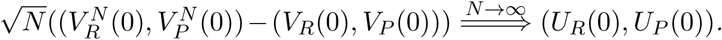 *Then, for the models* [*neu*], [*neg*] *and* [*pos*], 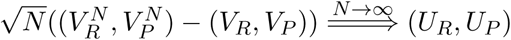, where (*U_R_*, *U_p_*) *solves*

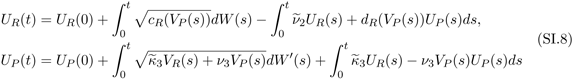

*with W, W′ independent Brownian motions,*

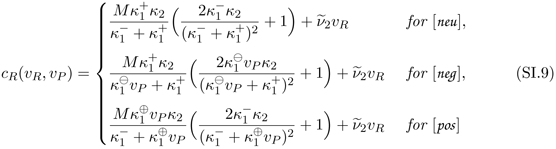

and

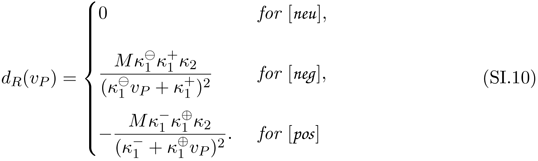

Remark C.1. In equilibrium, we have

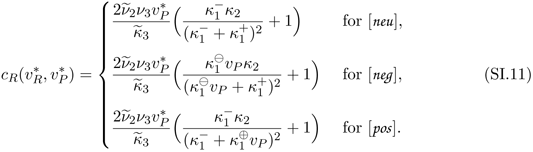

*Proof*. First, note that

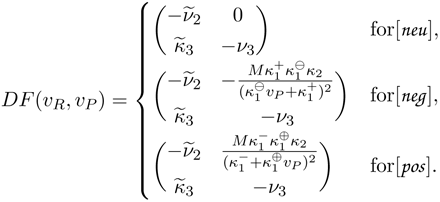

Again, we have to show (➊)-➍) from Section E in all cases for *F* as above, which already satisfies (➊). We focus on [*neu*] first. The system (*V*_on_,*V*_R_, *V*_P_) is a Markov process with a generator of the form (SI.21) with

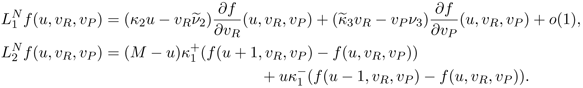

This implies that

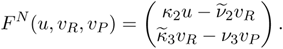

Hence, for (➊), we have to solve

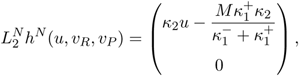

which is

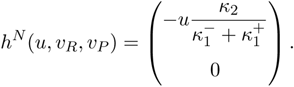

For the quadratic variation in (➍), we find that with *z^⊗2^ = zz^T^*

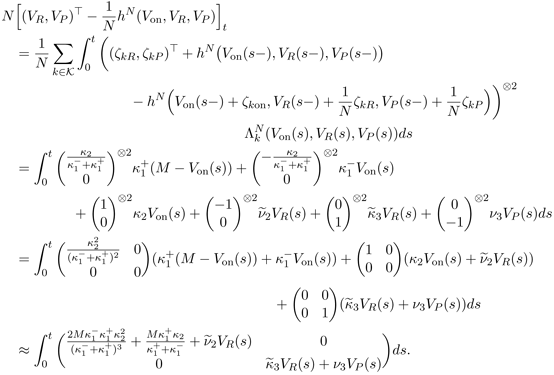

This shows the assertion for [*neu*]. The cases [*neg*] and [*pos*] are similar, if we change 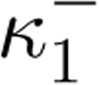 by 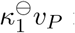 for [*neu*] and 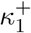 by 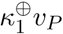 for [*pos*].□

### C.2 Equilibrium Fano factor…

Let us start in equilibrium, i.e. 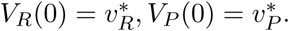. Then, we will plug in *c_R_* from (SI.11), and obtain Ornstein-Uhlenbeck processes in all cases. Since they are two-dimensional, their equilibrium (normal) distribution can be computed (see Section F).

…for [*neu*]

We obtain

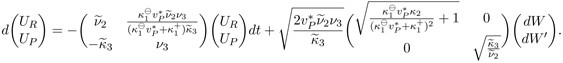

Hence, with *U* = (*U_R_*, *U_p_*)^T^, in equilibrium, from Corollary F.1,

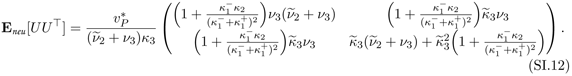

Therefore,

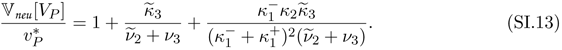

…for [*neg*]

Here, we obtain

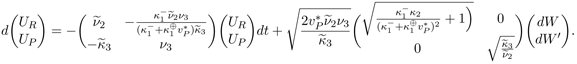

Hence, in equilibrium, and if…denote the quantities from (SI.12), and for 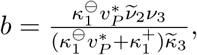,

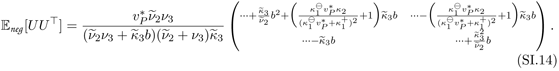

so

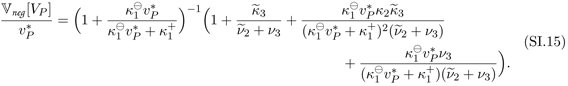

…for [*pos*]

We obtain

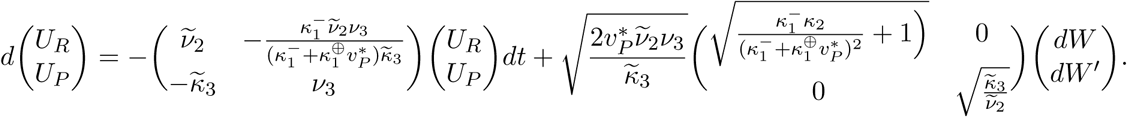

In equilibrium, we now have exactly (SI.14) but with 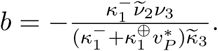. Hence,

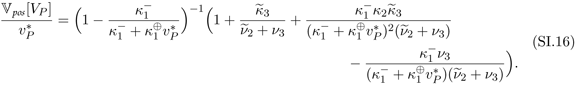

## D Comparison to previous results

### D.1 Comparison for [*neu*]

The neutral case is by now well-studied. From Paulsson (2005), we see that this equilibrium obeys, with

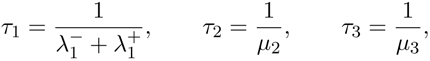

the expected life-times of a change in gene activity, mRNA and protein, respectively, that

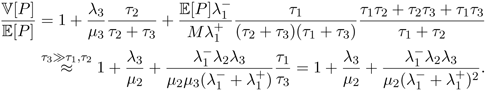

Since our scaling is exactly such that τ_3_ » τ_1_,τ_2_, i.e. the protein is much more stable than RNA and the state of the gene, this result is in line with [11] in the main text.

### D.2 Comparison with Thattai

In Thattai and van Oudenaarden (2001), a linearization of [*neg*] was studied in the case of fast switching on and off of the gene. (This will mean that both, 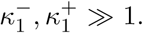) Consider as in the proof of Theorem 1 that

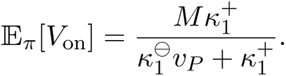

Then, study the system (compare with (●_neg_))

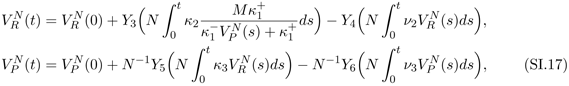

which can in the case (8) (i.e. 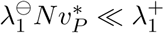) be approximated by using

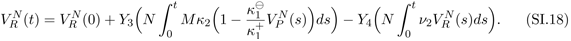

Now, the system (SI.17) and (SI.18) is exactly as on p. 3 in Thattai and van Oudenaarden (2001) with

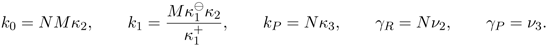

Plugging these variables into [3] of Thattai and van Oudenaarden (2001) we obtain in equilibrium with

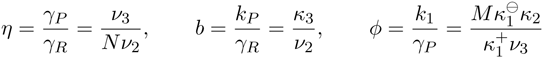

that (note that η is negligible since *N* is large and φ is small by (8))

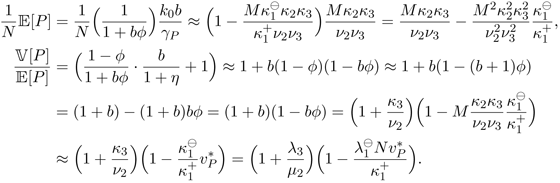

### D.3 Comparison with Swain

In Swain (2004) the author uses *d*_0_ = *μ*_2_,*d*_1_ = μ_3_,*v*_1_ = λ_3_,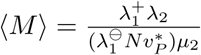 and

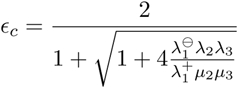

in order to obtain the Fano factor for the auto-regulatory gene expression with negative (transcriptional) feedback. Expressing terms as in Section C, he gets

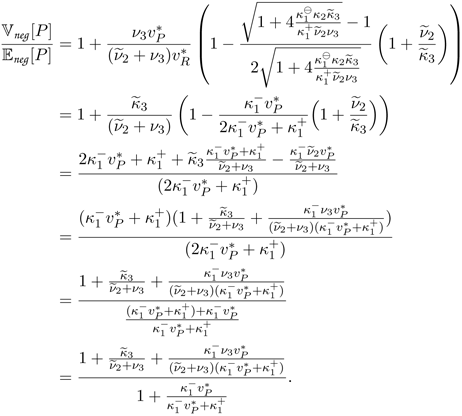

This is very similar to our equation (SI.15) except for a missing term emerging from gene expression. As explained in the main text, the Langevin approximation Swain uses cannot account for all fluctuations when a quasi-steady-state assumption is made. (Precisely, it cannot account for fluctuations in the averaged variables.)

### D.4 Simpler model of gene expression

We want to derive the simpler system without RNA from the full model. Noting that in (*_*neu*_), for κ_2_ = ν_2_ » 1, we have that for every finite *N*, RNA is approximately in equilibrium, which is Poisson with parameter 1 for every *on-gene*. Feeding this into the equation for *P*, we obtain that

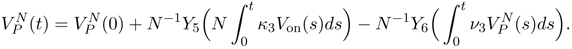

This is the same equation which is need for the simple model given in the main text. Hence, all results follow for *κ*_2_ = ν_2_ → ∞.

## E Recalling the approach of Kang et al. (2014)

We will consider a general system of chemical reactions with *S* as the set of chemical species and *K* the set of reactions; see also Anderson and Kurtz (2015) for further reference. The chemical reactions have the form

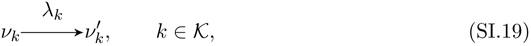

where 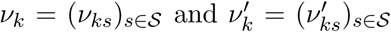 are vectors of chemical species, i.e. elements of 𝕅^S^. For the dynamics, we assume mass action kinetics, i.e. we set

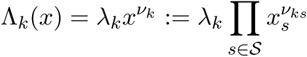

for the reaction rate of reaction *k* ∊*K*. With

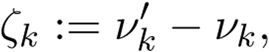

we can then define the dynamics of the Markov process *X* = (*X_s_*)_s∊S_ through the process *R_k_*, which describes the number of occurrences of reaction k up to time t. We have

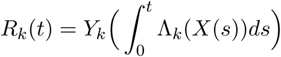

for independent unit rate Poisson processes *Y_k_*, *k* ∊ *K* and therefore

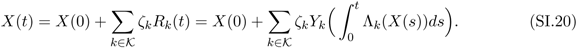

We will tailor the results of Kang et al. (2014) to the special case we need in our gene expression example. This means that we can make use of several simplifications, e.g. on the form of the generator of the full process.

1. Find *F*: 𝕉^ds^ such that

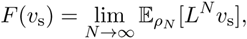

2 Find *h^N^* such that 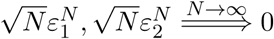 with 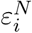 and 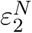 from (SI.24).

For some scaling parameter *N*, assume that 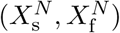 is a Markov jump process with state space 𝕅^ds^ x 𝕅^df^ such that for 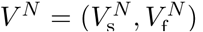 with 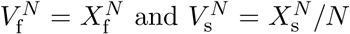, the system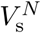 is a slow (rescaled) sub-system and 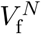 is a fast sub-system. We assume that the generator *L^N^* has the form

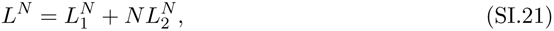

where 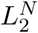 describes the dynamics of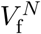, i.e. 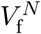 = 0 if *f* only depends on *v_s_*. Our goal is to show convergence

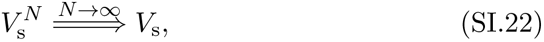

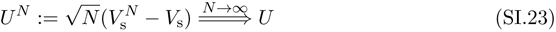

for some Vs and U. Therefore, we proceed as follows.

1. We have that (with the projection π_s_ on the slow species and 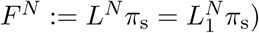)

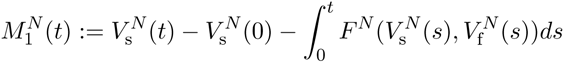

is a (local) martingale. For the convergence (SI.22), we assume that

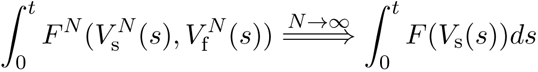

for some unique process V_s_, which holds for

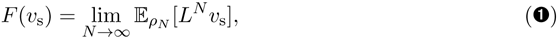

where ρ_N_ is the equilibrium of 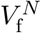 for fixed slow species v_s_. Thus, the convergence 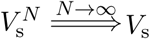 holds with

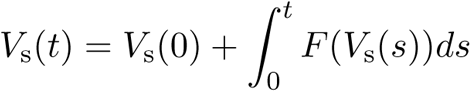

and we have shown (SI.22).

2. Note that

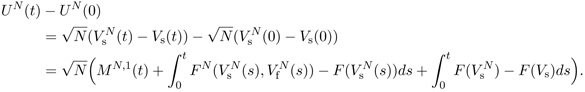

Assume that we

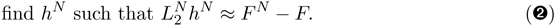

(The ‘≈’ is controlled by 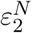 below. Note that this is a Poisson equation.) With

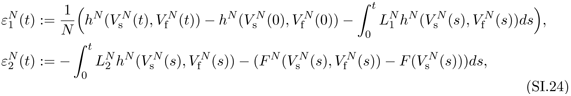

we obtain that

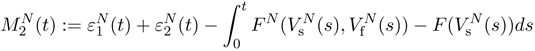

is a (local) martingale. Hence

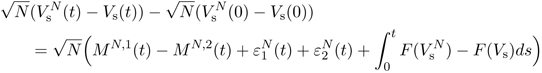

We assume for 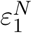,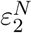 from (SI.24) that

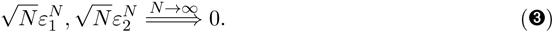

3. In order to show convergence of 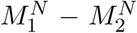, we use the Martingale (Central Limit Theorem; see Theorem 5. Note that since the quadratic variation of all integrals *f* dt vanishes, we find that (recall the notation for Chemical Reaction Networks from (SI.19)-(SI.20), which we now equip with a superscript *N* to account for the scaling constant), with z^⊗2^ = zz^T^

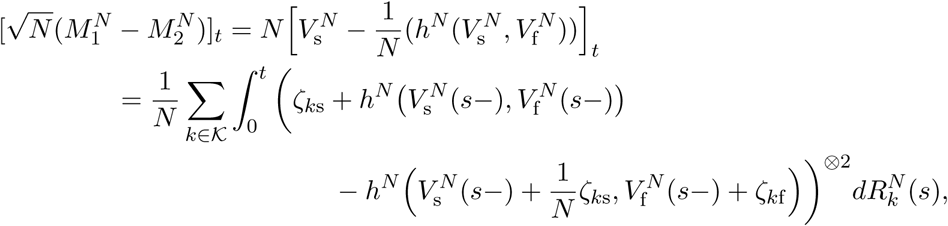

where ζ_*ks*_ and ζ_*kf*_ are the stochiometric changes of the *k*—th reaction in the slow and fast subsystem, respectively. Note that in all applications, we will have that 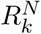 either changes slowly, or changes fast and can thus be approximated by a deterministic curve, such that

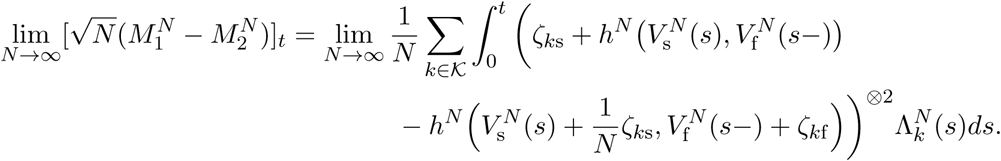

Now, for the equilibrium ρ_*N*_ of the fast species for given concentration of slow species, *v_s_*, if

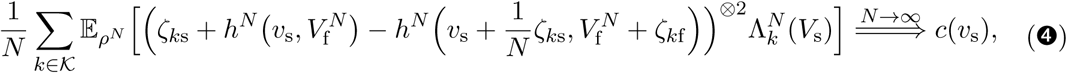

we have that

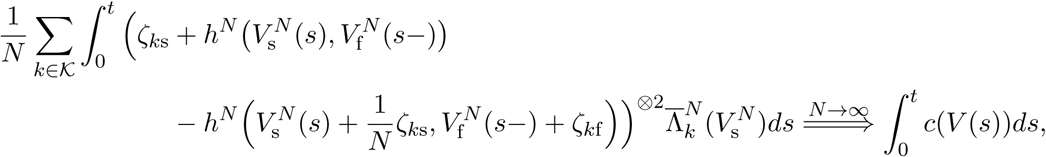

where the right hand side is a deterministic, absolutely continuous, 𝕉^s×s^-valued function with non-negative time-derivative. Hence we know from Theorem 5 that

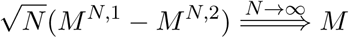

where *M* satisfies

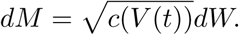

4. Concluding, if *F* ∊ C ^1^(𝕉^d_s_^) with

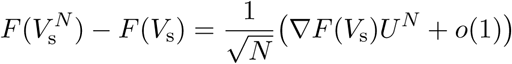

we find that, if 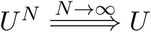 then

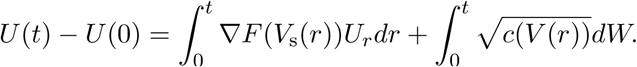

This gives (SI.23).

## F The two-dimensional Ornstein-Uhlenbeck process

We recall results for the two-dimensional Ornstein-Uhlenbeck process; see e.g. Gardiner (2009).

### Theorem 4 (Stationary variance of two-dimensional Ornstein-Uhlenbeck process)

*Let X* = (*X*_1_, *X*_2_) *solve*

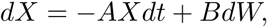

*where* A, B ∊ 𝕉^2×2^. *Then, if all eigenvalues of A have positive real part, the stationary solution X*_0_ *of the SDE has* 𝔼[*X*_0_] = 0 *and*

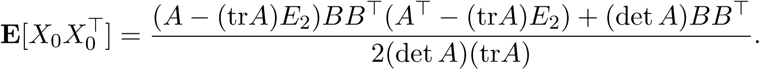

***Proof.*** Using partial integration, it is easy to see that this SDE is solved by

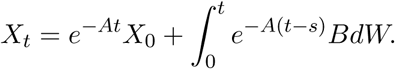

If all eigenvalues of A have positive real part, the stationary solution of the SDE has the distribution

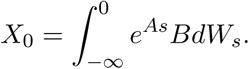

In particular, E[X_0_] = 0 and

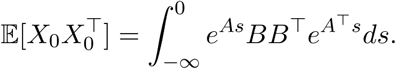

In order to compute the right hand side, we note that, for any 2 × 2-matrix 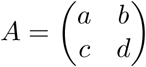, we have that (for the unit matrix *I*_2_)

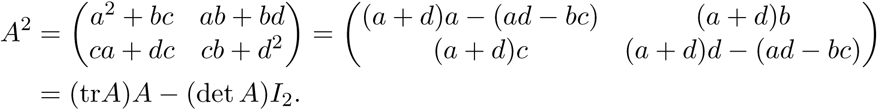

Hence, we can write

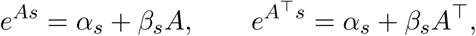

i.e

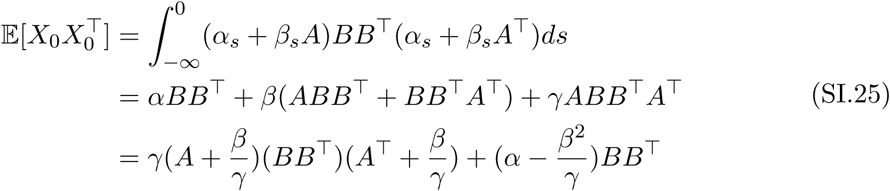

for

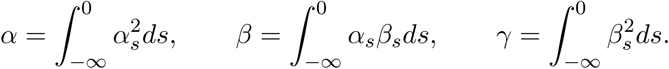

We then write

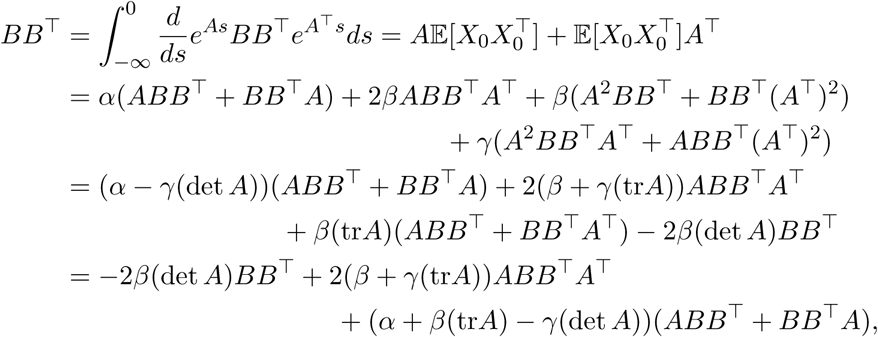

which is only possible if

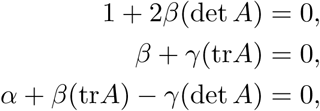

i.e

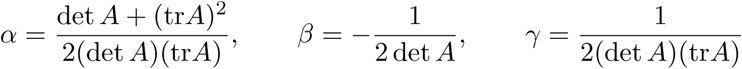

Combining this with (SI.25) then gives the result.□

**Corollary F.1** (Diagonal matrix B). ***Note that if***

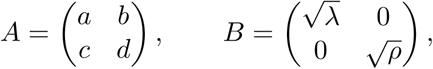

*then*

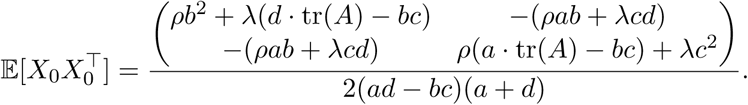

***Proof.*** Indeed, 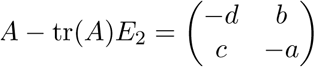 so

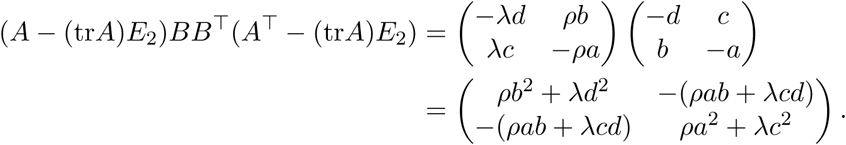
□

## G Martingale Central Limit Theorem

Here, we give a version of the martingale central limit theorem as given in Ethier and Kurtz (1986), Theorem 7.1.4.

### Theorem 5

*Let* (*M_N_*) *be a sequence of* 𝕉^d^*-valued martingales. Suppose that*

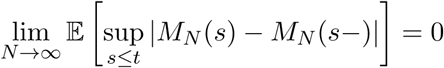

*and*

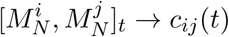

*for all t ≥ 0, where C = (c_ij_) is deterministic and continuous. Then* 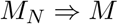*for a stochastic process M which is Gaussian with independent increments and* 𝔼[*M*(*t*)*M*(*t*)^T^] = *C*(*t*).

Furthermore, since *C* is non-negative and symmetric and if *C* is absolutely continuous its derivative is non-negative and thus has a non-negative square-root. This implies that *M* can be written as

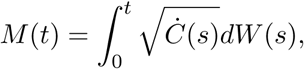

where *W* is the *d*-dimensional standard Brownian motion.

## References

[1] Swain PS, Elowitz MB, Siggia ED. Intrinsic and extrinsic contributions to stochasticity in gene expression. Proc Natl Acad Sci USA. 2002;99(20):12795-12800.

[2] Elowitz MB, Levine AJ, Siggia ED, Swain PS. Stochastic gene expression in a single cell. Science. 2002;297(5584):1183-1186.

[3] Raj A, A van Oudenaarden A. Nature, nurture, or chance: stochastic gene expression and its consequences. Cell. 2008;135(2):216-226.

[4] Li GW, Xie XS. Central dogma at the single-molecule level in living cells. Nature. 2011;475:308-315.

[5] Raser JM, O’Shea EK. Noise in gene expression: origins, consequences, and control. Science. 2005;309(5743):2010-2013.

[6] Singh A, Soltani M. Quantifying intrinsic and extrinsic variability in stochastic gene expression models. PloS one. 2013;8(12):e84301.

[7] Singh A. Transient changes in intercellular protein variability identify sources of noise in gene expression. Biophysical journal. 2014;107(9):2214-2220.

[8] Kaern M, Elston TC, Blake WJ, Collins JJ. Stochasticity in gene expression: from theories to phenotypes. Nat Rev Genet. 2005;6(6):451-464.

[9] Maamar H, Raj A, Dubnau D. Noise in gene expression determines cell fate in Bacillus subtilis. Science. 2007;317(5837):526-529.

[10] Eldar A, Elowitz MB. Functional roles for noise in genetic circuits. Nature. 2010;467(7312):167-173.

[11] Silva-Rocha R, de Lorenzo V. Noise and robustness in prokaryotic regulatory networks. Annual review of microbiology. 2010;64:257-275.

[12] Wang Z, Zhang J. Impact of gene expression noise on organismal fitness and the efficacy of natural selection. Proc Natl Acad Sci. 2011;108(16):E67-E76.

[13] Paulsson J. Models of stochastic gene expression. Physics of Life Review. 2005;2:157-175.

[14] Kumar N, Singh A, Kulkarni RV. Transcriptional Bursting in Gene Expression: Analytical Results for General Stochastic Models. PLoS computational biology. 2015;11(10):e1004292.

[15] Lestas I, Vinnicombe G, Paulsson J. Fundamental limits on the suppression of molecular fluctuations. Nature. 2010;467(7312):174-178.

[16] Hornung G, Barkai N. Noise propagation and signaling sensitivity in biological networks: a role for positive feedback. PLoS computational biology. 2008;4(1):e8.

[17] Thattai M, van Oudenaarden A. Intrinsic noise in gene regulatory networks. Proc Nat Acad Sci. 2001;98(15):8614-8619.

[18] Swain PS. Efficient Attenuation of Stochasticity in Gene Expression Through Post-transcriptional Control. Journal of Molecular Biology. 2004;344(4):965-976.

[19] Dessalles R, Fromion V, Robert P. A Stochastic Analysis of Autoregulation of Gene Expression. Preprint. 2016;p. 1-27.

[20] Ethier SN, Kurtz TG. Markov processes: characterization and convergence. Wiley series in probability and mathematical statistics. New York, Chichester: J. Wiley & Sons; 1986.

[21] Kurtz T. Solutions of ordinary differential equations as limits of pure jump Markov processes. Journal of Applied Probability. 1970;7:49-58.

[22] Darling RWR. Fluid Limits of Pure Jump Markov Processes: a Practical Guide. arxiv preprint. 2002;https://arxiv.org/abs/math/0210109:1-16.

[23] Kurtz T. Limit Theorems for sequences of jump Markov processes approximating ordinary differential processes. Journal of Applied Probability. 1970;8:344-356.

[24] Seegel LA, Slemrod M. The Quasi-Steady-State Assumption: a Case Study in Perturbation. SIAM Review. 1989;31(3):446-477.

[25] Ball K, Kurtz TG, Popovic L, Rempala G. Asymptotic analysis of multiscale approximations to reaction networks. Ann Appl Probab. 2006;16(4):1925-1961.

[26] Kang HW, Kurtz T, Popovic L. Central Limit Theorems and Diffusion Approximations for Multiscale Markov Chain Models. Ann Appl Probab. 2014;24:721-759.

[27] Bokes P, King JR, Wood A, Loose M. Multiscale stochastic modelling of gene expression. J Math Biol. 2012;65(3):493-520.

[28] Pardoux E, Veretennikov AY. On Poisson equation and diffusion approximation 1. Ann Probab. 2001;29:1061-1085.

[29] Pardoux E, Veretennikov AY. On Poisson equation and diffusion approximation 2. Ann Prob. 2003;31:1166-1192.

[30] Bar-Even A, Paulsson J, Maheshri N, Carmi M, O’Shea E, Pilpel Y, et al. Noise in protein expression scales with natural protein abundance. Nature genetics. 2006;38(6):636-643.

[31] Gillespie D. Exact stochastic simulation of coupled chemical reactions. J Phys Chem. 1977;81:2340-2361.

[32] Hornos JEM, Schultz D, Innocentini G, Wang J, Walczak A, Onuchic J, et al. Self-regulating gene: an exact solution. Physical review E, Statistical, nonlinear, and soft matter physics. 2005;72(5 Pt 1):051907.

[33] Shahrezaei V, Swain PS. Analytical distributions for stochastic gene expression. Proc Natl Acad Sci USA. 2008;105(45):17256-17261.

[34] Ramos A, Innocentini G, Hornos J. Exact time-dependent solutions for a self-regulating gene. Physical review E, Statistical, nonlinear, and soft matter physics. 2011;83(6 Pt 1):062902.

[35] Grönlund A, Lötstedt P, Elf J. Transcription factor binding kinetics constrain noise suppres-sion via negative feedback. Nature communications. 2013;4:1864.

[36] Kim JK, Josić K, Bennett MR. The relationship between stochastic and deterministic quasisteady state approximations. BMC systems biology. 2015;9:87.

[37] Becskei A, Serrano L. Engineering stability in gene networks by autoregulation. Nature. 2000;405(6786):590-593.

[38] Cardelli L, Csikász-Nagy A, Dalchau N, Tribastone M, Tschaikowski M. Noise Reduction in Complex Biological Switches. Scientific reports. 2016;6:20214.

[39] Munsky B, Neuert G, van Oudenaarden A. Using gene expression noise to understand gene regulation. Science. 2012;336(6078):183-187.

## References

Anderson, D. and T. G. Kurtz (2015). Stochastic Analysis of Biochemical Systems. Springer.

Ball, K., T. G. Kurtz, L. Popovic, and G. Rempala (2006). Asymptotic analysis of multiscale approximations to reaction networks. Ann. Appl. Probab. 16(4), 1925-1961.

Dessalles, R., V. Fromion, and P. Robert (2016). A stochastic analysis of autoregulation of gene expression. Preprint, 1-27.

Ethier, S. N. and T. G. Kurtz (1986). Markov processes: characterization and convergence. Wiley series in probability and mathematical statistics. New York, Chichester: J. Wiley & Sons.

Gardiner, C. (2009). Stochastic Methods. A Handbook for the Natural and Social Sciences. Springer.

Kang, H.-W., T. Kurtz, and L. Popovic (2014). Central limit theorems and diffusion approximations for multiscale Markov chain models. Ann. Appl. Probab. 24, 721-759.

Paulsson, J. (2005). Models of stochastic gene expression. Physics of Life Review 2, 157-175.

Swain, P. S. (2004). Efficient attenuation of stochasticity in gene expression through post-transcriptional control. Journal of Molecular Biology 344 (4), 965-976.

Thattai, M. and A. van Oudenaarden (2001). Intrinsic noise in gene regulatory networks. Proc. Nat. Acad. Sci. 98 (15), 8614-8619.

